# The contributions from the progenitor genomes of the mesopolyploid Brassiceae are evolutionarily distinct but functionally compatible

**DOI:** 10.1101/2020.08.10.245258

**Authors:** Yue Hao, Makenzie E. Mabry, Patrick P. Edger, Michael Freeling, Chunfang Zheng, Lingling Jin, Robert VanBuren, Marivi Colle, Hong An, R. Shawn Abrahams, Jacob D. Washburn, Xinshuai Qi, Kerrie Barry, Christopher Daum, Shengqiang Shu, Jeremy Schmutz, David Sankoff, Michael S. Barker, Eric Lyons, J. Chris Pires, Gavin C. Conant

## Abstract

The members of the tribe Brassiceae share a whole genome triplication (WGT), and one proposed model for its formation is a “two-step” pair of hybridizations producing hexaploid descendants. However, evidence for this model is incomplete, and the evolutionary and functional constraints that drove evolution after the hexaploidy are even less understood. Here we report a new genome sequence of *Crambe hispanica*, a species sister to most sequenced Brassiceae. Using this new genome and three others that share the hexaploidy, we traced the history of gene loss after the WGT using POInT (the Polyploidy Orthology Inference Tool). We confirm the two-step formation model and infer that there was a significant temporal gap between those two allopolyploidizations, with about a third of the gene losses from the first two subgenomes occurring prior to the arrival of the third. We also, for the 90,000 individual genes in our study, make parental “subgenome” assignments, inferring, with measured uncertainty, which of the progenitor genomes of the allohexaploidy each gene derives from. We further show that each subgenome has a statistically distinguishable rate of homoeolog losses. There is little indication of functional distinction between the three subgenomes: the individual subgenomes show no patterns of functional enrichment, no excess of shared protein-protein or metabolic interactions between their members, and no biases in their likelihood of having experienced a recent selective sweep. We propose a “mix and match” model of allopolyploidy, where subgenome origin drives homoeolog loss propensities but where genes from different subgenomes function together without difficulty.

## Introduction

Fifty years ago, Ohno (Ohno 1970) published his forceful opus on the role of gene duplication, and in particular of *genome* duplication (aka polyploidy), in evolutionary innovation. Since then, evidence both of polyploidy’s ubiquity (Wolfe and Shields 1997; Soltis and Soltis 2012; Van de Peer et al. 2009, 2017) and of its role in evolutionary innovations such as yeast aerobic glucose fermentation, the organization of the retinae of teleost fishes and in plant defensive compounds has continued to accumulate (Conant and Wolfe 2007; Merico et al. 2007; van Hoek and Hogeweg 2009; Edger et al. 2015; Sukeena et al. 2016). Preeminent among the polyploid lineages are the flowering plants, where over 180 ancient polyploidies are known (One Thousand Plant Transcriptomes Initiative 2019).

When a new polyploid genome is created by the merging of similar but not identical progenitor species, it is referred to as an allopolyploid. Among allopolyploidies, the preferential retention of gene copies (homoeologs) from one of the parental subgenomes, known as biased fractionation, has been observed in yeast, maize, cotton, monkeyflower, *Arabidopsis*, *Brassica*, and nematodes (Thomas et al. 2006; Conant and Wolfe 2008a; Cheng et al. 2012; Parkin et al. 2014; Renny-Byfield et al. 2015; Edger et al. 2017; Emery et al. 2018; Schoonmaker et al. 2020). Allopolyploids also show a tendency for genes from one of the subgenomes to be more highly expressed, and silencing or loss of genes from the remaining subgenomes is correspondingly more likely (Thomas et al. 2006; Schnable et al. 2011; Yoo et al. 2014). A number of sources of these biases have been proposed, from variations in transposon silencing (Freeling et al. 2012; Woodhouse et al. 2014; Zhao et al. 2017; Alger and Edger 2020), to the disruption of organelle-nucleus communication (Sharbrough et al. 2017; Costello et al. 2019) and epigenetic changes due to the genomic shock of polyploidy (McClintock 1984; Wendel et al. 2018; Bird et al. 2018). In this work, we sought to critically evaluate one such proposal: that allopolyploids might bring together coevolved and conflicting copies of multi-protein complexes (Codoñer and Fares 2008; Gong et al. 2012; Scienski et al. 2015; Emery et al. 2018). In this framework, early random gene losses from one subgenome that partly resolved these conflicts might then set the polyploidy down a path favoring losses from that subgenome. A related proposal was made by Makino and McLysaght (2012), who argued that selection to maintain dosage balance among interacting genomic neighbors could produce local, and eventually global, biases in fractionation.

It is also notable that not all homoeologs are equally likely to revert to single-copy after a polyploidy, regardless of the level of biased fractionation. Duplicated genes coding for transcription factors, ribosomal proteins and kinases are over-retained after independent polyploidies in flowering plants, yeasts, ciliates and vertebrates (Seoighe and Wolfe 1998; Blanc and Wolfe 2004; Maere et al. 2005; Aury et al. 2006; Makino and McLysaght 2010). These patterns are best explained by a need to maintain dosage balance among highly interacting genes (Birchler et al. 2005; Hakes et al. 2007; Birchler and Veitia 2012, 2014; Conant et al. 2014). There are also genes that prefer *not* to be duplicated: genes for DNA repair and those targeted to organelles have returned to single-copy rapidly after genome duplication (De Smet et al. 2013; Conant 2014).

The Brassiceae are the most morphologically diverse tribe in the family Brassicaceae (Cheng et al. 2014) and contain important crops such as broccoli, cabbage, kale, mustard and canola. This tribe experienced a hexaploidy (a.k.a. whole genome triplication; WGT) between 5 and 9 MYA, after its divergence from *Arabidopsis thaliana* (Wang et al. 2011). This Brassiceae WGT is a valuable system for studying all of the phenomena mentioned above because the triplication allows us to explore each in unusual detail. This polyploidy was originally inferred with comparative linkage mapping (Lagercrantz 1998; Lukens et al. 2004; Parkin et al. 2005; Schranz et al. 2006) and confirmed by chromosome painting (Lysak et al. 2005; Lysak 2009). The patterns of biased fractionation observed in the genome of *Brassica rapa* suggested that the triplication “event” was actually two separate allopolyploid hybridizations involving three distinct diploid progenitor species, with the merger of the two currently highly fractionated ancestral subgenomes occurring first, followed by the subsequent addition of a third subgenome, which currently possesses the most retained genes (Tang et al. 2012; Cheng et al. 2012). However, this proposal is worth revisiting as it rests on inferences from a single genome: a phylogenetically broader analysis of the genomes that descend from the hexaploidy would more firmly ground our descriptions of its early history. At the moment, we lack genomes from early-diverging lineages with the hexaploidy, such as those in the genus *Crambe*, which is sister to the genus *Brassica* (Arias and Pires 2012). Biologically, *Crambe* species are not only important industrial oilseed sources because of their high erucic acid content (Lazzeri et al. 1997; Warwick and Gugel 2003; Carlsson et al. 2007) but also could serve as resources for *Brassica* crop development (Rudloff and Wang 2011).

Using a new genome sequence from *Crambe hispanica*, we analyzed the Brassiceae WGT with our tool for modeling post-polyploidy genome evolution: POInT (the Polyploidy Orthology Inference Tool; Conant and Wolfe 2008a). We sought to first confirm the two-step hexaploidy model and its relationship to the observed three subgenomes in the extant genomes. POInT, which we recently extended to allow the analysis of WGTs (Schoonmaker et al. 2020), is ideally suited to this task, because it can model homoeolog losses phylogenetically and test for biases in fractionation without *ad hoc* assumptions. We then tested the proposal that functional differences between the allopolyploid progenitors contributed to the biases in homoeolog losses using functional hierarchies, gene co-expression information, protein interaction catalogs and metabolic network data.

## Results

### *A well-assembled and annotated genome of* Crambe hispanica

The genome of *Crambe hispanica* was assembled using PacBio reads. This assembly had a contig N50 of 4.4 Mb across 1,019 contigs with a total assembly length of 480 Mb. Eleven terminal telomeres were resolved by the Canu assembler (Koren et al. 2017). The assembly graph showed low heterozygosity and few assembly artifacts, with the exception of one mega-cluster consisting of a high copy number LTR across 500 contigs and spanning ~30 Mb. The draft assembly was then polished using Illumina paired-end data. We also used Hi-C proximity ligation sequencing data to scaffold the genome, which resulted in 18 scaffolds that include 99.5% of the original assembly with a scaffold N50 of 32.6 Mbp and scaffold N90 of 30.1 Mbp. The annotated genome is of high quality: we compared its gene set against the Benchmarking Universal Single-Copy Orthologs (BUSCO v.2; Simão et al. 2015) plant dataset (embryophyta_odb9), finding that 95.8% of these expected genes were present in our annotation.

### Inferring blocks of triple-conserved synteny in four triplicated Brassiceae genomes and estimating an ancestral gene order

Based on their phylogenetic placement and assembly quality, we selected and retrieved from CoGe (Lyons and Freeling 2008; Lyons et al. 2008a) three additional mesohexaploid genomes for our analyses: those of *Brassica rapa* (version 1.5, CoGe id 24668; Wang et al. 2011), *Brassica oleracea* (TO1000 version 2.1, CoGe id 26018; Liu et al. 2014; Parkin et al. 2014) and *Sinapis alba* (version 1.1, CoGe id 33284). For each of these four genomes, we inferred blocks of triple conserved synteny (TCS), with the genome of *Arabidopsis thaliana* used as an unduplicated reference. We then merged these blocks across all of the four genomes: we refer to each such locus as a “pillar”. Each pillar consists of between 1 and 3 surviving genes in each of the four genomes. As described in the *Methods*, we used both a set of TCS blocks inferred with POInT containing 14,050 pillars (*P_pillars_*) and a separate ancestral genome reconstruction that estimates the gene order that existed just prior to the WGT. The latter contains five reconstructed ancestral chromosomes involving 89 scaffolds with a total of 10,868 ancestral genes. When we match these genes to the TCS blocks computed with POInT, the result is 7,993 ancestrally-ordered pillars (*A*_pillars_).

### Inferring the evolutionary relationships of the four Brassiceae genomes from gene loss patterns

We fit models of WGT evolution (see below) to several different orderings of the 14,050 pillars in the *P_pillars_* set and to the *A_pillars_* (Supplemental Table S1). These orderings of the *P_pillars_* differed in their number of synteny breaks: we used the ordering with the highest likelihood under the WGT 3rate G1Dom model for our remaining analyses (see below). Similarly, we compared the fit of three possible phylogenetic topologies to the pillars under this model: the remainder of our analyses use the topology shown in Figure 1, which has the highest likelihood. We note that one of the other two topologies, while having a lower likelihood under POInT’s models (Supplemental Fig S1), is the phylogeny estimated using the plastid genome (Arias and Pires 2012). Because the *A_pillars_* give similar parameter estimates but comprise a smaller dataset, we will discuss our results in terms of the *P_pillars_*.

**Figure 1.**
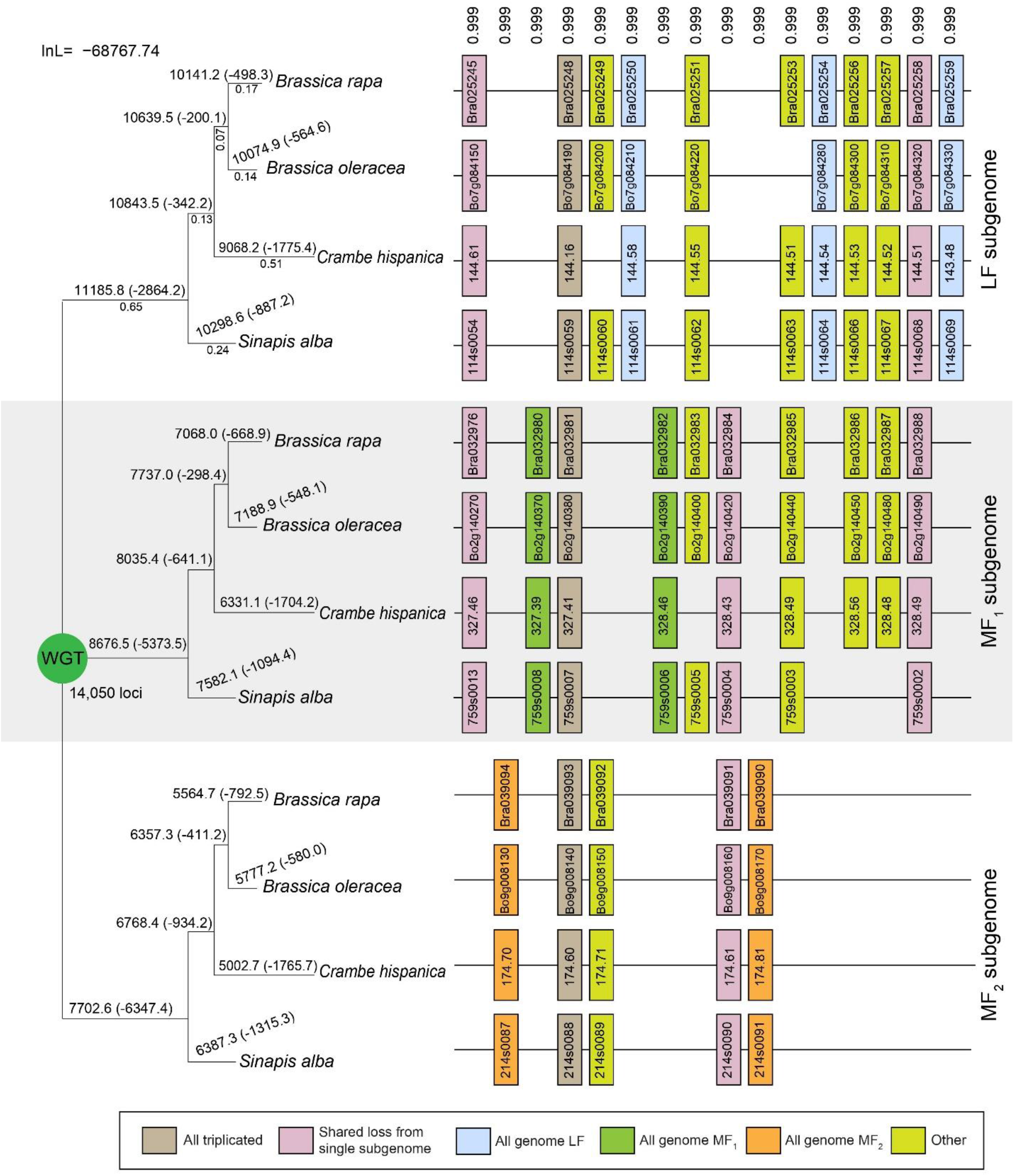
Subgenome assignment and inference of gene loss after the shared WGT in four species. After the WGT, each ancestral locus could potentially expand to three gene copies, but due to biases in the loss events, the number of surviving genes from the subgenomes are unequal. Our analyses (see *Results*) indicate the presence of a less fractionated (LF) subgenome and two more fractionated ones (MF1 and MF2). These inferences are based on the gene losses observed across four genomes and along the phylogeny depicted. Shown here is a window of 16 post-WGT loci (out of the total 14,050 such loci) in four species that share the WGT: *Brassica rapa*, *Brassica oleracea*, *Crambe hispanica* and *Sinapis alba*. Each pillar corresponds to an ancestral locus, and the boxes represent extant genes. Pairs of genes are connected by lines if they are genomic neighbors (e.g., in synteny). The numbers on top of each pillar are the posterior probabilities assigned to this combination of orthology relationships relative to the other *(3!)^4^−1=1295* possible orthology states. The numbers above each branch of the tree give the number of genes in each subgenome surviving to that point, with the number of gene losses in parentheses. The gene loss inferences made by POInT are probabilistic: as some gene losses cannot be definitively assigned to a single branch, the resulting loss estimates are not integers. The numbers below the branches in the first subtree are POInT’s branch length estimates (αt).

### The three subgenomes differ in their propensity for homoeolog copy loss

POInT employs user-defined phylogenetic Markov models of gene loss after WGT. These models have seven states (Figure 2): the triplicated state **T** in which all three copies from the WGT are still present; the “duplicated” states **D_1,2_**, **D_1,3_**, **D_2,3_** where one out of the three gene copies has been lost, and three single-copy state **S_1_**, **S_2_**, and **S_3_**. Previous work suggested that the three subgenomes that formed these hexaploids are distinct in their patterns of gene preservation (Tang et al. 2012; Cheng et al. 2012), consisting of a “less fractionated” genome (LF), a subgenome with intermediate levels of gene loss (more fractionated 1 or MF1) and an even more fractionated subgenome (MF2). We hence defined state **S_1_** to correspond to LF and **S_2_** and **S_3_** to MF1 and MF2, respectively.

**Figure 2.**
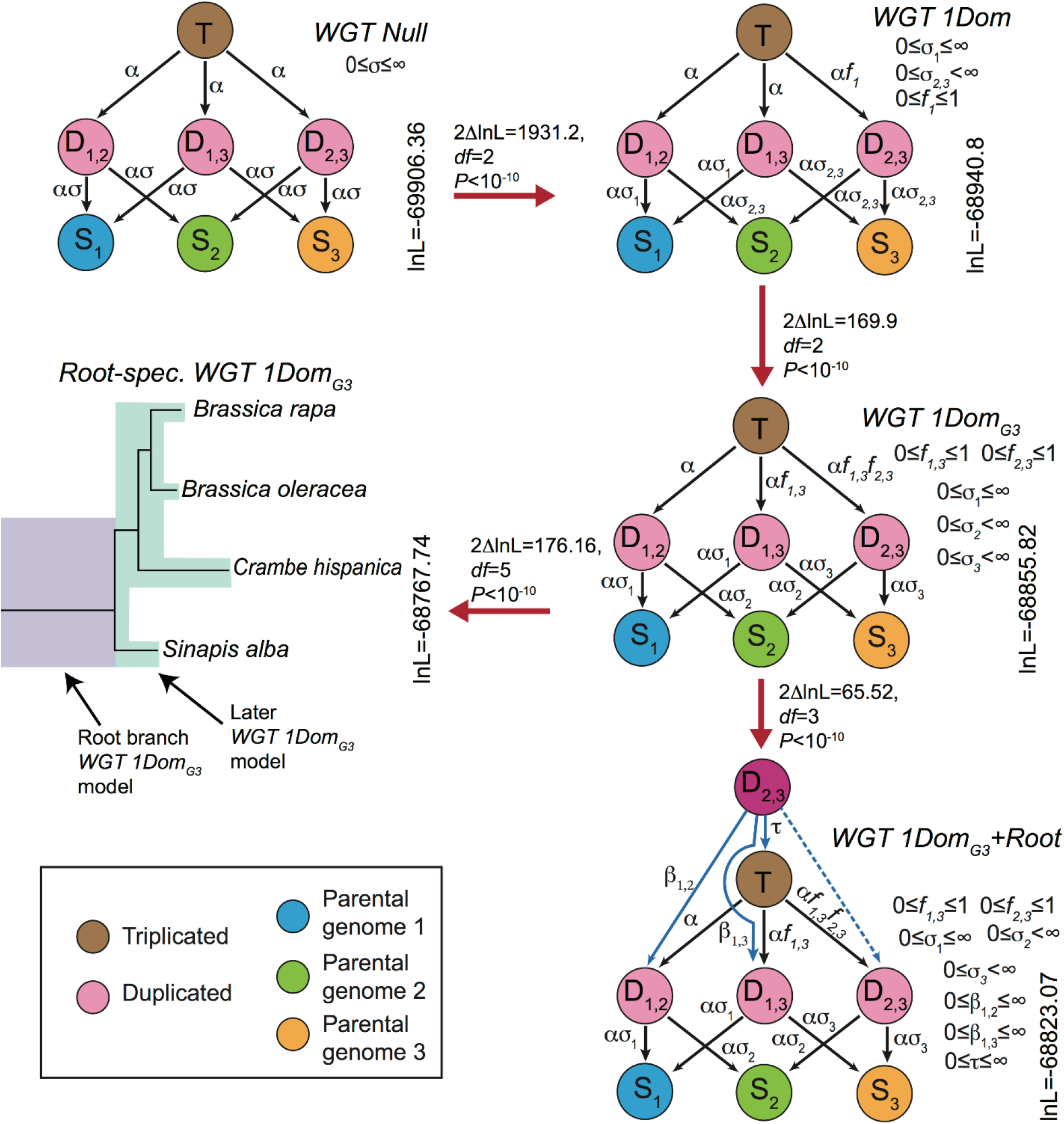
POInT’s models for inferring WGT. Five different models of post-WGT evolution and their ln-likelihoods are shown. In each model, the colored circles represent different states. The brown circle represents the triplicated state (**T**); the pink circles are duplicated states (**D_1,2_**, **D_1,3_** and **D_2,3_**); the blue, green and yellow circles are three single-copy states (**S_1_** for the LF subgenome, **S_2_** for the MF1 subgenome and **S_3_** for the MF2 subgenome). The transition rates between states are shown above the arrows. α: transition rate from triplicated state to duplicated states; ασ: transition rates from duplicated states to single copy states; f: fractionation parameters; β and τ: root model parameters. Red arrows connect pairs of models compared using likelihood ratio tests (see *Methods*). *WGT Null model*: transition rates are the same across three subgenomes, modeling the scenario of no biased fractionation. *WGT 1Dom model*: with the biased fractionation parameter f_1_ (0 ≤ f_1_ ≤ 1), the MF1 and MF2 subgenomes are more fractionated than LF subgenome. *WGT 1Dom_G3_ model*: two fractionation parameters f_1,3_ and f_2,3_ were introduced, distinguishing the three subgenomes: MF2 is more fractionated than MF1, and MF1 is more fractionated than LF. *Root-spec*. *WGT 1Dom_G3_ model*: similar to the previous model, but with two sets of parameters, one set for the root branch and the other for the remainder of the branches. *WGT 1Dom_G3_ + Root model*: Two-step hexaploidy model created by starting each pillar in an intermediate state **D_2,3_**. This state represents the merging of the MF1 and MF2 subgenomes as the first step of the hexaploid formation. The **T, D_1,2_,** and **D_1,3_** states represent the second allopolyploidy, with either no prior homoeolog losses (**T**) or a loss from one of the two MF subgenomes prior to that event (**D_1,2_,** or **D_1,3_**).

POInT statistically assigned genes from each of the four mesopolyploid genomes to the LF, MF1 and MF2 subgenomes with high confidence: 75% of the pillars have subgenome assignments with posterior probabilities > 0.84 (Supplemental Fig S3). We observe clear signals of biased fractionation: while we estimate that 2,864 genes were lost from the LF subgenome along the shared root branch (e.g., prior to the split of *S. alba* from the other three species), the corresponding figures for MF1 and MF2 are 5,373 and 6,347 respectively (Figure 1). These values are in qualitative agreement with previous findings (Xie et al. 2019; Liu et al. 2014; Cheng et al. 2014, 2012).

We assessed the statistical support for these estimated differences in the subgenomes’ rates of homoeolog loss using a set of nested models of post-WGT gene loss. We started with a model (WGT Null) that did not differentiate between the subgenomes, meaning that the shared base transition rate from **T** to **D_1,2_**, **D_1,3_** or **D_2,3_** is defined to be α (0 ≤ α < ∞, Figure 2). The transition rate from **D_1,2_**, **D_1,3_** or **D_2,3_** to **S_1_**, **S_2_** or **S_3_** is scaled by σ: e.g., occurs at rate α×σ. We compared this model to a more complex one that allowed losses of both triplicated and duplicated genes to be less frequent from a posited less-fractionated subgenome LF (WGT 1Dom, Figure 2). This model introduces a fractionation parameter *f_1_* (0 ≤ *f_1_* ≤ 1), which potentially makes the transitions between T and D_2,3_ rarer than the other T-to-D rates (α×f_1_; see Figure 2). The WGT 1Dom model fits the pillar data significantly better than does WGT Null (Figure 2; *P<*10^−10^, likelihood ratio test with two degrees of freedom). We next compared the WGT 1Dom model to a WGT 1Dom_G3_ model that gives MF1 and MF2 separate loss rates. Again, this model gives a better fit to the pillar data than did WGT 1Dom (*P*<10^−10^, likelihood ratio test with two degrees of freedom, Figure 2). We hence confirm the presence of three subgenomes, distinguishable by their patterns of homoeolog loss. It is important to recall here that our approach does not require the identification of these three subgenomes *a priori*: the probabilistic assignment of genes to subgenomes is an integral part of the POInT orthology computation: as a result, the inherent uncertainty in these assignments is accounted for in estimating the various biased fractionation parameters. Our orthology inferences can be explored visually with POInT_browse_ (http://wgd.statgen.ncsu.edu/).

### Patterns of post-WGT gene loss support the “two-step” model of hexaploidy

To test the hypothesis that the WGT proceeded in two steps (Cheng et al. 2012; Tang et al. 2012), we used two approaches. First, we applied an extended version of the WGT 1Dom_G3_ model where each model parameter was allowed to take on distinct values on the root branch and on the remaining branches (Root-spec. WGT 1Dom_G3_ in Figure 2). This extended model fits the pillar data significantly better than does the original WGT 1Dom_G3_ model (*P*<10^−10^, likelihood ratio test with five degrees of freedom, Figure 2). The biased fractionation parameters for the root branch differ from those of the remaining branches: the value of f_1,3_ on the root is smaller than on later branches (0.6445 versus 0.7368) while f_2,3_ is larger (0.6766 versus 0.4078). These values are consistent with a two-step hypothesis: prior to the arrival of LF, there would have been a number of losses from MF1 and MF2, meaning that the relative preference for LF would be higher (smaller f_1,3_).

In our second approach, we developed a specific model of the two-step hexaploidy (WGT 1Dom_G3_+Root_LF_ in Figure 2). This model describes the transition from a genome *duplication* to a triplication: all pillars start in state **D_2,3_**: e.g., the first allopolyploidy has just occurred and the MF1 and MF2 genes are present but not the LF ones. We then model the addition of LF as transitions to either the **T**, **D_1,2_** or the **D_1,3_** states (with rates τ, β_1,2_ or β_1,3_, respectively). State **T** is seen when no losses occurred prior to the arrival of LF, the other states occur when either MF1 or MF2 experienced a loss prior to the arrival of LF. Any pillars that remain in **D_2,3_** had no corresponding gene arrive from LF. Of course, at the level of the individual pillar, we have insufficient data to make such inferences: the utility of this model is to give global estimates of the degree of fractionation seen in MF1 and MF2 prior to the arrival of LF. This model offers a significantly improved fit over WGT 1Dom_G3_ (*P<*10^−10^, likelihood ratio test with three degrees of freedom, Figure 2). More importantly, we can propose other versions of this model where either MF1 or MF2 is the last arriving subgenome: when we do so, the model fit is much worse than seen with WGT 1Dom_G3_+Root_LF_ model (Supplemental Table S1). Hence, we can conclude that subgenomes MF1 and MF2 had already begun a process of (biased) fractionation prior to the addition of the LF subgenome. Note that these conclusions derive only from genes that were inferred to be present in all three parental subgenomes, a requirement of the POInT models.

### A gap between the two allopolyploidies

This root-specific model also allows us to estimate the state of MF1 and MF2 immediately before the arrival of LF. In particular, we can estimate the percentage of pillars that had already experienced losses prior to LF’s arrival. About 28% of all of the MF1 homoeologs inferred to have been lost on the root branch were lost prior to the arrival of LF, with the equivalent number of MF2 losses being 38%. A negligible 0.3% of pillars do not appear to have received a copy of the LF homoeolog.

### Mixed evidence for differences in selective constraint between subgenomes

In our dataset there 218 loci that have retained triplicates in all four genomes and have subgenome assignment confidence ≥ 95%. For each loci, we calculated the selective constraints the group of 12 genes using codeml (Yang 2007), allowing the genes from each subgenome to have a different *d_N_/d_S_* value. On average, among these retained triplets, genes from the LF subgenome show slightly smaller *d_N_/d_S_* values than do those from MF1 and MF2, but these differences are not statistically significant (Wilcoxon rank sum tests LF - MF1: *P* = 0.300, LF - MF2: *P* = 0.079; Supplemental Fig S4).

### Single copy genes from multiple subgenomes are enriched in genes functioning in DNA repair

GO overrepresentation tests were performed with the *Arabidopsis* orthologs of genes returned to single copy by the end of the root branch from each subgenome. Similar to previous findings (De Smet et al. 2013), we found that single copy genes are enriched in biological processes such as DNA repair and DNA metabolism (Supplemental Fig S5). More specifically, single copy genes from the LF subgenome are enriched in base-excision repair, while MF1 single copy genes are enriched in nucleotide-excision repair, non-recombinational repair and double-strand break repair (Supplemental Fig S5A). Single copy genes from both LF and MF1 show overrepresented molecular functions in endo- and exodeoxyribonuclease activities (Supplemental Fig S5B). LF single copy genes are also enriched in RNA interference processes, suggesting that such interference, targeted to the MF1 and MF2 subgenomes, could be one mechanism by which biased fractionation was driven.

### Genes from the same subgenome are not overly likely to physically or metabolically interact

For genes with high subgenome assignment confidence (≥ 95%), we mapped those assignments (LF, MF1 or MF2) and the duplication status at the end of the root branch onto the nodes (gene products) of the *A. thaliana* protein-protein interaction (PPI) network (*Methods*). For comparative purposes, we also produced a mapping of an extant network, based on the gene presence/absence data and subgenome assignments in *B. rapa*. In the “ancient” network inferred at the end of the common root branch, there are a relatively large number of nodes (1,952) associated with surviving triplicated loci: these nodes were connected by a total of 2,384 triplet-to-triplet edges. The *B. rapa*-specific network contains fewer nodes with retained triplets (662): there were 263 edges connecting these nodes (Figure 3A).

**Figure 3.**
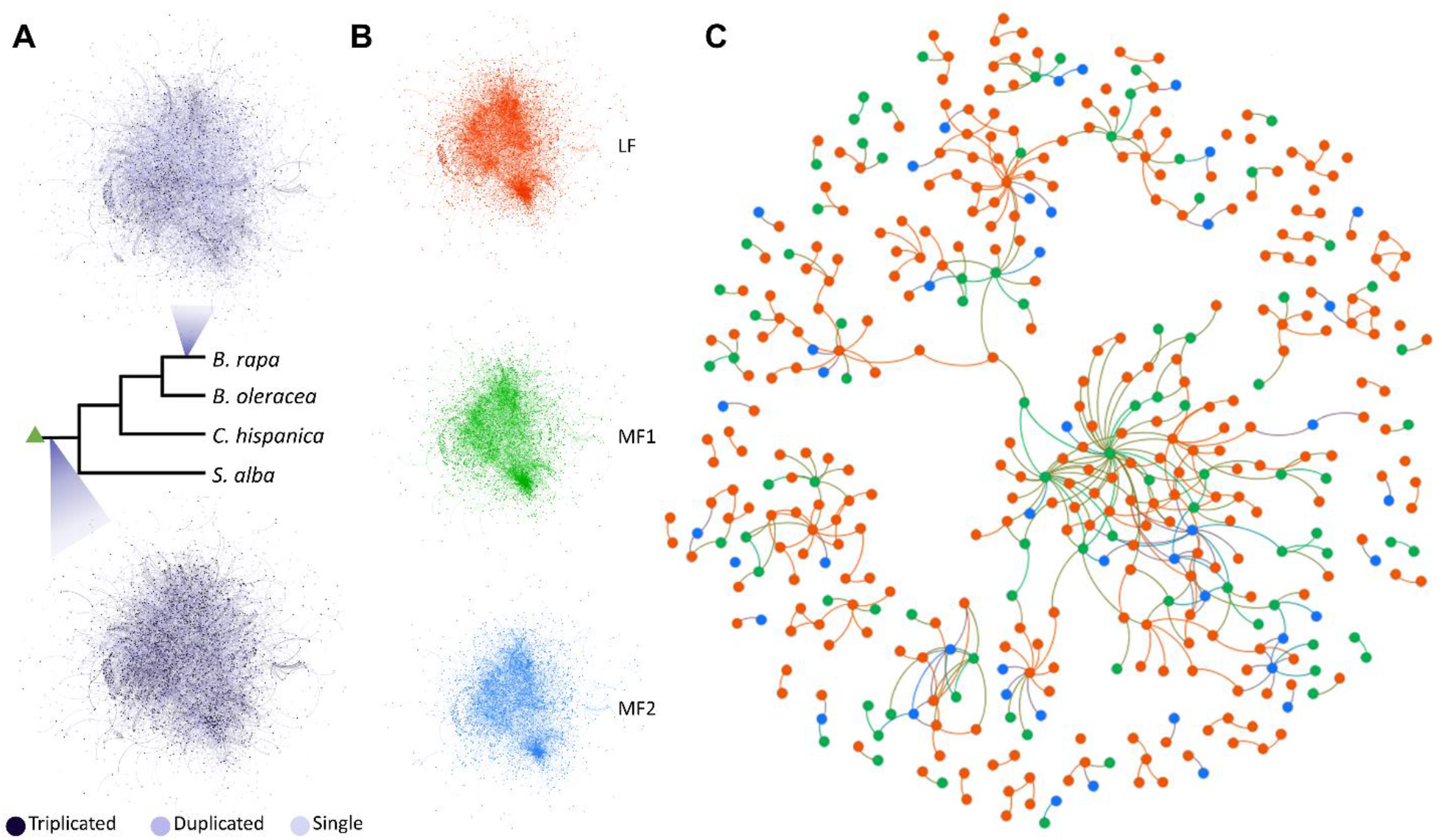
Protein-protein interaction networks after the WGT. (*A*) The *Arabidopsis* PPI network at the root branch (bottom) and the same PPI network colored by the *Brassica* rapa gene retention status (top). The dark purple nodes represent retained triplets. See Supplemental Code. (*B*) the PPI network partitioned by subgenome assignment at the root branch. LF subgenome: red - 4,249 nodes and 8,454 edges. MF1 subgenome: green - 3,379 nodes and 6,442 edges. MF2 subgenome: blue - 3,073 nodes and 4,961 edges. (*C*) A subset of the PPI network where only nodes encoded by single copies genes and connected to other single copy nodes are shown. Red nodes are from the LF subgenome, green nodes are from the MF1 subgenome and blue nodes are from the MF2 subgenome.

The dosage constraints that affect surviving gene copies post-polyploidy will tend to result in the retention of genes involved in multiunit complexes or in the same signaling pathways (Birchler and Veitia 2007, 2012; Conant et al. 2014). Thus, we expected to see that the retained triplets showed higher network connectivity. And indeed, our permutation tests reveal that the retained triplets on the root branch are significantly over-connected to each other in the PPI network (*P* = 0.018, Supplemental Fig S6). We also hypothesized that proteins coded for from the same subgenome would be more likely to be connected due to preferential retention of genes from a single complex from the same subgenome. To test this idea, we partitioned the gene products based on their subgenome of origin. The LF subgenome contains more genes and thus more exclusive connections (Figure 3B). When considering only genes that had returned to single-copy by the end of the root (Figure 3C), we identified 188 LF-LF edges among 886 single copy LF genes, with fewer edges exclusive to MF1 and MF2 genes (30 and 3, respectively). We used randomization (see *Methods*) to test whether the numbers of such subgenome-specific edges differed from what would be expected by chance. When considering the network as a whole, we found that there were significantly *fewer* LF-LF edges than expected (*P* = 0.022; Supplemental Fig S6). However, when we considered only the single-copy genes in the network, the number of subgenome-specific edges did not differ from that seen in random networks for any of the three subgenomes (*P* = 0.286 for LF-LF edges, see Supplemental Fig S6), suggesting that the original dearth of such edges was a statistical artifact resulting from the excess of triplet-to-triplet edges.

We also explored the association of between genes’ role in metabolism and their pattern of post-hexaploidy evolution using the *A. thaliana* metabolic network (*Methods*). However, again considering the state of each pillar at the end of the root branch, we did not find an excess of shared metabolic interactions between triplicated or single-copy genes in this network (Supplemental Fig S6).

Finally, we asked whether genes from the same subgenome are more likely to be co-expressed. We constructed a *B. rapa* co-expression network from the RNA-seq data described in the *Methods* section. In this network edges connect pairs of genes that are highly correlated in their expression (*Methods*). The inferred co-expression network contains 3,933 nodes (e.g., genes) from the LF subgenome, 2,310 nodes from MF1 and 1,982 from MF2. We then counted the number of edges connecting pairs of nodes from the same subgenome. To assess whether there was an excess of such shared subgenome co-expression relationships, we randomly rewired the network 100 times and compared the edge count distributions from these randomized networks to those of the real network (Pérez-Bercoff et al. 2011). We found that the real network did not show a significant excess of shared edges between genes from the same subgenome when compared to the randomized networks (LF-LF *P*=0.36, MF1-MF1 *P*=0.82, MF2-MF2 *P*=0.08, Figure 4A-F).

**Figure 4.**
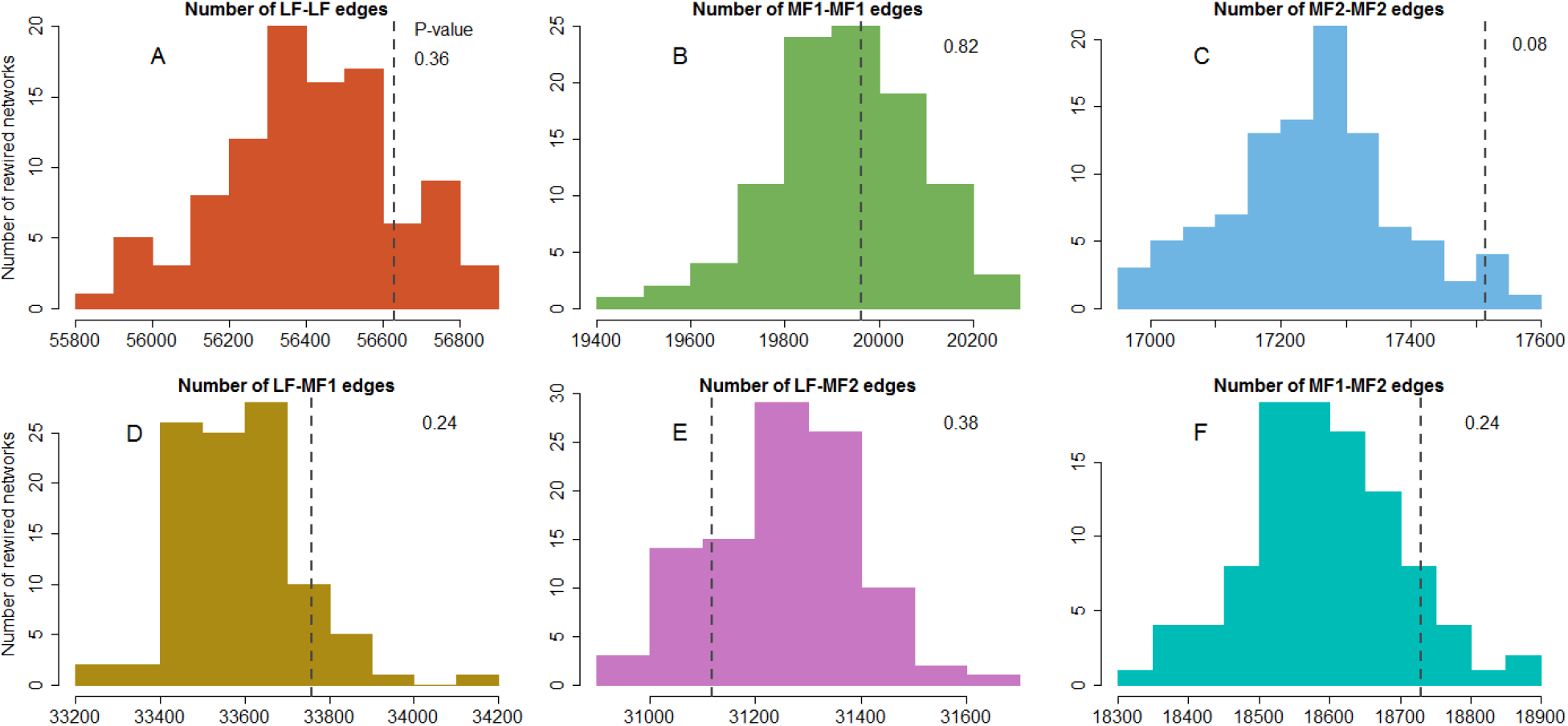
Subgenome-specific edge counts for 100 rewired *Brassica rapa* co-expression networks compared to those from the actual network. (*A*) Distribution of the number of edges connecting pairs of *B. rapa* genes both from the LF subgenome in 100 rewired networks. (*B*) Distribution of the number of edges connecting pairs of genes both from the MF1 subgenome. (*C*) Distribution of the number of edges connecting pairs of genes both from the MF2 subgenome. (*D*) Distribution of the number of edges connecting LF genes to MF1 genes. *(E)* Distribution of the number of edges connecting LF genes to MF2 genes. (*F*) Distribution of the number of edges connecting MF1 and MF2 genes. In each panel, the dark grey dashed line shows the number of edges with that set of subgenomes assignments for the true network. See Supplemental Code.

### Subgenome of origin does not affect the propensity to have experienced a selective sweep

We tested for associations between genes’ subgenome of origin and their propensity to experience recent selective sweeps. Data on these sweeps was taken from a recent scan in *B. rapa* by Qi et al. (2021). No subgenome had either an excess or a deficit of observed sweeps relative to the other two (Supplemental Fig S7). Genes from the MF1 subgenome showed slightly negative association with selective sweeps (*P* = 0.0089, chi-square test).

## Discussion

The combination of the new genome sequence of *Crambe hispanica* and our modeling of the post-WGT evolution of the four Brassiceae genomes using POInT allowed us to draw a number of conclusions regarding the Brassiceae WGT. We confirmed previous work (Tang et al. 2012; Cheng et al. 2012) arguing that these genomes derive from a pair of ancient allopolyploidies: more subtly, we also show that, as had been proposed, the least fractionated subgenome (e.g., the one with the most retained genes) is very likely the genome that was added last. To these proposals, we add the new observation that these hybridization events were likely not particularly closely spaced in time: our model predicts that on the order of 1/3 of the gene losses from subgenomes MF1 and MF2 that occurred on the root branch occurred *before* the arrival of the LF subgenome. Of course, one should not take this result to necessarily imply a very large number of calendar years between the events: gene loss immediately after polyploidy can be quite rapid (Scannell et al. 2007; De Smet et al. 2013). In the future, it will be interesting to further refine the timing of these events: the problem, however, is a challenging one because the allopolyploid nature of the events means that molecular clock approaches will tend to estimate speciation times for the allopolyploid ancestors rather than hybridization times.

Many forces shape genome evolution after polyploidy. A tendency for genes that operate in multiunit complexes or involved in signaling cascades to remain duplicated post-polyploidy is best explained by the presence of dosage constraints driven by a need to maintain the stoichiometry and kinetics of assembly for such functional units (Birchler et al. 2005; Birchler and Veitia 2007, 2012; Conant et al. 2014; Birchler et al. 2016). On the other hand, genes involved in functions such as DNA repair very often return rapidly to singleton status after duplication (Freeling 2009; De Smet et al. 2013). Our results illustrate the importance of these dosage effects, with genes whose products interact with many other gene products in *A. thaliana* being overly likely to be retained in triplicate in these Brassicae genomes. This pattern is not observed for metabolic genes, a result we interpret as illustrating metabolism’s dynamic robustness to gene dosage changes (Kacser and Burns 1981).

We had previously argued that one force driving the biased fractionation that distinguishes the LF, MF1 and MF2 subgenomes might be selection to maintain coadapted complexes from a single parental subgenome (Emery et al. 2018). That such coadapted complexes exist and respond to polyploidy is suggested by the gene conversions seen after the yeast polyploidy among the duplicated ribosomal and histone proteins (Evangelisti and Conant 2010; Scienski et al. 2015). However, these examples may be exceptions rather than the rule, meaning that pressure to maintain coadapted complexes is not a significant driver of biases in fractionation. We found that although there was some degree of functional distinction for single copy genes from the LF subgenome (e.g., enrichment in biological processes such as DNA repair and RNA interference), more generally speaking, there was no significant evidence of functional incompatibilities between single-copy genes from different subgenomes. Thus, genes from the same subgenome were not more likely to interact with each other physically, nor were the genes returned to single copy on the common root branch functionally subdivided among the subgenomes. And even the DNA repair enzyme genes that rapidly returned to single-copy appear to derive from at least two of the three subgenomes. It hence appears that De Smet et al.’s (2013) original hypothesis that these genes may be prone to dominant negative interactions may best explain their preference for a single-copy state.

It remains to be seen if the “mix and match” pattern of subgenome retention observed here represents the dominant mode of evolution for allopolyploidies. Of course, whether or not subgenome conflicts exist may be partly a question of the preexisting differences between the progenitor species, and a more general survey of allopolyploidies that includes estimates of the progenitor genomes’ divergence prior to the polyploidy events would be most enlightening. If the pattern holds, however, the implications would be significant, as hybridization represents an important means of adaption (Paterson 2005; Hollister 2015; Alix et al. 2017; Blanc-Mathieu et al. 2017; Smukowski Heil et al. 2017). Adding the effects of hybridization to polyploidy’s known association with innovation (Edger et al. 2015) and to the tendency of dosage sensitive genes to remain duplicated for the longer periods needed for such innovations (Blanc and Wolfe 2004; Conant and Wolfe 2008b; Conant et al. 2014; Zhao et al. 2017; Liang and Schnable 2018; Qiu et al. 2020) makes a strong case for considering polyploidy a critical source of material for innovation at the genomic level.

## Methods

### Crambe hispanica *(PI 388853) sample preparation, genome sequencing*

Leaf tissue was harvested from 36 dark treated inbred plants (selfed for nine generations; PI 388853). Dark treatment was performed to reduce chloroplast abundance and involved leaving the plants in a dark room for 3-4 days. After treatment, 5g of tissue was collected across 36 plants. This process was repeated three times, allowing us to obtain a total of 15g of tissue. This tissue was then sent to the University of Delaware Sequencing and Genotyping Center at the Delaware Biotechnology Institute (Newark, DE, USA) for high molecular weight DNA isolation and library preparation prior to PacBio (Pacific Biosciences, Menlo Park, CA, USA) and Illumina (San Diego, CA) sequencing. Libraries were prepared using standard SMRTbell procedures, followed by sequencing of 11 PacBio SMRT cells on a PacBio sequel and one PacBio SMRT cell of RSII sequencing. Paired-end 150 bp reads were generated on an Illumina HiSeq 2500 system. For Hi-C scaffolding, 0.5g tissue sample was sent to Phase Genomics (Phase Genomics Inc. Seattle, WA, USA).

### Crambe hispanica *v1.1 genome assembly and annotation*

The assembly of the *Crambe hispanica* v1.1 genome was performed using Canu v1.6 (Koren et al. 2017). In total, 3.9 million raw PacBio reads spanning 48 Gb were used as input for Canu. The following parameters were modified for assembly: minReadLength=1000, GenomeSize=500Mb, corOutCoverage=200 “batOptions=-dg 3 -db 3 – dr 1 -ca 500 -cp 50”. All other parameters were left as default. The assembly graph was visualized using Bandage (Wick et al. 2015) to assess ambiguities in the graph related to repetitive elements and heterozygosity. The draft Canu assembly was polished reiteratively using high-coverage Illumina paired-end data (82 million reads) with Pilon v1.22 (Walker et al. 2014). Quality filtered Illumina reads were aligned to the genome using Bowtie 2 v2.3.0 (Langmead and Salzberg 2012) under default parameters and the resulting BAM file was used as input for Pilon with the following parameters: --flank 7, --K 49, and --mindepth 8. Pilon was run recursively three times using the updated reference each time to correct the maximum number of residual errors.

A Proximo Hi-C library was prepared as described (Phase Genomics Inc. Seattle, WA, USA) and sequenced on an Illumina HiSeq 2500 system with paired-end 150 bp reads. The *de novo* genome assembly of Hi-C library reads were used as input data for the Phase Genomics Proximo Hi-C genome scaffolding platform.

The genome was annotated using MAKER (Campbell et al. 2014), using evidence of protein sequences downloaded from the Araport 11 and Phytozome 12 plant databases (Cheng et al. 2017; Goodstein et al. 2012) and *C. hispanica* transcriptome data. The transcriptome data for genome annotation was extracted from bud, root, and leaf tissues under standard daylight conditions using the Thermo Fisher Scientific PureLink RNA Mini Kit. Library prep was done using Illumina TruSeq DNA PCR-free and sequenced for non-stranded mRNA-Seq 2×250 on Illumina HiSeq. *C. hispanica* transcriptomic data were assembled with StringTie (Pertea et al. 2015). Repetitive regions in the genome were masked using a custom repeat library and Repbase Update (Bao et al. 2015) through RepeatMasker Open-4.0 (Smit et al. 2015). *Ab initio* gene prediction was performed using SNAP (Korf 2004) and AUGUSTUS (Stanke and Waack 2003). The resulting MAKER gene set was filtered to select gene models with Pfam domain and annotation edit distance (AED) < 1.0. Then, the amino acid sequences of predicted genes were searched against a transposase database using BLASTP and an E-value cutoff of 10^−10^ (Campbell et al. 2014). If more than 30% of a given gene aligned to transposases after the removal of low complexity regions, that gene was removed from the gene set.

### Triple-conserved synteny reconstruction

We have developed a three-step pipeline for inferring the conserved synteny blocks created by polyploidy (Emery et al. 2018). For the first step of this pipeline, we used *Arabidopsis thaliana* Col-0 version 10.29 (CoGe genome id 20342) as a nonhexaploid outgroup and identified homologous genes between it and each of the four hexaploid genomes using GenomeHistory (Conant and Wagner 2002). Genes were defined as homologous if their translated products shared 70% percent amino acid sequence identity and the shorter sequence was at least 80% percent of the length of the longer. In the second step, we sought to place genes from each of the hexaploid genomes into blocks of triple-conserved synteny (TCS) relative to their *A. thaliana* homologs. To do so, we inferred a set of “pillars,” each of which contains a single gene (or group of tandem duplicates) from *A. thaliana* and between 1 and 3 genes from the hexaploidy genome. Using simulated annealing (Kirkpatrick et al. 1983; Conant and Wolfe 2006), we sought a combination of pillar gene assignments and relative pillar order that maximized the TCS. In the third and final step, we merged the pillars across the four hexaploid genomes, using their *A. thaliana* homologs as indices. We then sought a global pillar order that minimized the number of synteny breaks across all of the hexaploid genomes (Supplemental Fig S2). These three steps resulted in a set of 14,050 ordered pillars, each with at least one surviving gene from each of the four genomes (Figure 1) and a corresponding “ancestral” gene from *A. thaliana*. Supplemental Table S1 shows that POInT’s model inferences are consistent across a number of such estimated ancestral orders.

### An ancestral genome order reconstruction

As a verification of our POInT pipeline, we also sought an independent inference of the order of the genes in the parental subgenomes just prior to the first step of the *Brassica* triplication. First, we used CoGe’s SynMap (Lyons et al. 2008b) to identify homologs between the *A. thaliana* and *Arabidopsis lyrata* genomes and those between *B. rapa* and *B. oleracea*. The SynMap algorithm was applied with a chaining distance of 50 genes and a minimum of five aligned gene pairs to identify likely orthologous genes in all pairwise-comparisons of the four genomes. Paralogs were identified by self-comparisons of each of the two *Brassica* genomes with SynMap. Then these orthologs and paralogs were grouped into 24,011 homology sets with the ‘OMG!’ program (Zheng et al. 2011). Every homology set consists of 1-3 *Brassica* paralogs from each of the three *Brassica* genomes and a single *Arabidopsis* gene from each of the two *Arabidopsis* genomes, representing one “candidate gene” in the reconstructed ancestral genome. Among these, 2,178 homology sets contained the maximum of eight genes (one each from the two *Arabidopsis* genomes and three each from the two *Brassica* genomes).

The homology sets were used to retrieve the ancestral gene order from adjacency graph using an efficient algorithm called Maximum Weight Matching (MWM; Zheng et al. 2013). We identified all the gene adjacencies in the four genomes, considering only the genes in the homology sets. Each adjacency was then weighted according to how many of the 8 possible adjacencies were actually observed. The MWM produced an optimal set of 10,944 linear contigs containing all 24,001 putative ancestral genes from the homology sets that included 13,057 of 45,982 total adjacencies in the data set, with the remaining adjacencies being inconsistent with this optimal set. We used the contigs in the output of the MWM to reconstruct each of the 5 ancestral chromosomes. There were 34 contigs containing large proportions of genes originating in two or more of the ancient chromosomes that were discarded, as were any contigs containing four or fewer genes from a *Brassica* genome. While the 9,712 contigs so omitted represent 89% of all contigs, they represent only 55% of the genes, leaving a small group of large contigs with strong synteny relations in our ancestral reconstruction. We next identified adjacencies among the contigs themselves and applied a second iteration of MWM on them, giving the optimal ordering of those contigs. Combining these orders with the existing gene order information within each contig yields the position of all the genes on each ancestral chromosome. This order was mapped to our set of pillars of TCS, giving a subset of those pillars ordered by this ancestral order estimate.

### The phylogenetic relationships of the triplicated members of the Brassicaceae

POInT fits the models shown in Figure 2 to the pillar data under an assumed phylogenetic topology using maximum likelihood, allowing us to use that likelihood statistic to compare different phylogenetic relationships among these four hexaploid taxa. POInT’s computational demands were too great to allow testing all 15 rooted topologies of 4 species (POInT’s models are not time reversible). However, by making the reasonable assumption that *B. rapa* and *B. oleracea* are sister to each other, we were able to test the three potential relationships of *C. hispanica* and *S. alba* to the two *Brassicas*. Figure 1 gives the maximum likelihood topology: the two alterative topologies and their likelihoods are given in Supplemental Fig S1.

### Selective constraints of the retained triplets

We identified 218 pillars that retained triplicated genes across all four genomes and where the confidence in their subgenome assignments was ≥ 95%. For each such pillar, the 12 sequences were aligned using T-coffee (Notredame et al. 2000). The cladogram for each such set of 12 genes consists of three subtrees grouping four sequences that belong to same subgenome in the same sister group (Supplemental Fig S4). Using codeml in PAML (Yang 2007) with CodonFreq set to F3×4, we inferred three distinct *d_N_/d_S_* ratios: one for each of the three subtrees deriving from the three parental subgenomes.

### Functional analysis of single-copy genes from different subgenomes

We performed functional analysis for genes where we have high (≥ 95%) confidence that they returned to single copy along the common root branch. Using the corresponding “ancestral” locus from *A. thaliana*, we performed individual Gene Ontology analyses with PANTHER (Mi et al. 2019) Overrepresentation Tests (release date 20190711) for genes from each subgenome. The background list used in all cases was the loci that remained duplicated or triplicated at the end of the root branch.

### Protein-protein interaction and metabolic network analysis

The *A. thaliana* protein-protein interaction (PPI) network was downloaded from BioGRID (Stark et al. 2011; Arabidopsis Interactome Mapping Consortium 2011). The root-branch post-WGT subgenome assignments for each “ancestral” locus represented by an *Arabidopsis* gene were mapped onto the nodes (gene products) of the PPI network, so long as our confidence in those subgenome assignments was ≥ 95%. Similarly, for the extant *B. rapa*, we took loci with high subgenome assignment confidence ≥ 95% and mapped their *A. thaliana* orthologs onto network nodes. The resulting PPI network (Figure 3) was visualized using Gephi 0.9.2 (Bastian et al. 2009) with the Fruchterman Reingold and Yifan Hu layout algorithms (Fruchterman and Reingold 1991; Hu 2006). To test whether gene products from the same subgenome are over-connected in this network, we permutated the subgenome assignments 1,000 times, holding the network topology unchanged. We then compared the actual number of edges connecting single copy genes from the same subgenome with the distribution of this value seen in the randomized networks (Supplemental Fig S6). We also asked whether the ancestral genes corresponding to retained triplets showed an excess of connections amongst themselves. Because the number of edges between retained triplets and between single-copy genes are not independent, we performed an additional set of permutations, in which we held all the triplet rows constant and only shuffled the subgenome assignments of the remaining nodes.

We performed similar analyses using the AraGEM v1.2 metabolic network from *A. thaliana* (de Oliveira Dal’Molin et al. 2010; Bekaert et al. 2012). In this network, each node represents a biochemical reaction, and pairs of nodes are connected by edges if their respective reactions share a metabolite. For each *A. thaliana* gene encoding an enzyme catalyzing one such reaction, we mapped the root-branch subgenome assignments (again requiring ≥ 95% confidence), assigning to that gene three presence/absence variables (one per subgenome). Then, for each subgenome, we counted the number of edges between pairs of nodes with at least one pair of single-copy genes from a common subgenome. We assessed significance by holding the network topology and *Arabidopsis* gene assignments constant and randomizing the subgenome assignments 1,000 times. We then compared the distributions of the single-subgenome edge counts from the simulations with the actual values (Supplemental Fig S6).

### Brassica rapa *co-expression network analysis*

We generated a gene expression dataset for *Brassica rapa* spanning diverse experimental conditions, including the following: a cold treatment in leaves (4hrs and 28hrs post), methyl jasmonate treatment in leaves (4hrs and 28hrs post), anaerobic treatment in leaves (4 and 8hrs post), salt treatment in roots (4hrs and 28hrs post) and a diurnal time course in leaves (every 4hrs, 6 timepoints) in standard light-dark conditions but also in complete dark and complete light conditions. Total RNA was extracted from above organs using the Invitrogen Purelink RNA Mini Kit (Carlsbad, CA, USA), converted into a library using the Illumina TruSeq RNA kit (San Diego, CA, USA), and paired-end 100bp reads were sequenced on the HiSeq 2000 instrument at the VJC Genomics Sequencing Laboratory at the University of California, Berkeley. The NextGENe V2.17 (SoftGenetics, State College, PA, USA) software package was used to remove low-quality Illumina data, map reads to the *B. rapa* FPsc (v1.0, CoGe id 20101) reference genome, and calculate normalized RPKM (reads per kilobase of transcript per million) values for all genes.

We filtered the dataset to only include genes that were missing a measured expression value for at most one of the 32 RNA-seq libraries, leaving 24,907 *B. rapa* genes in it. The gene identifiers used for the expression dataset were from the *B. rapa* FPsc (v1.0, CoGe id 20101) reference genome, so we translated these identifiers to those from *B. rapa* Chiifu (v1.5, id 24668) using CoGe SynMap (Lyons et al. 2008b). In so doing, we only used *B. rapa* genes with one-to-one matches between the two references. For any pair of genes in the expression dataset, we calculated the Spearman’s correlation coefficient of their RPKM values. A co-expression network was then constructed using highly correlated gene pairs, e.g., pairs having Spearman’s correlation coefficients ≥ 0.9 (positive correlations), or ≤ −0.9 (negative correlations). Thus, the nodes of this co-expression network are *B. rapa* genes, and the edges represent correlation in expression. The co-expression network was randomized 100 times by rewiring the edges, while holding the nodes and their subgenome assignments unchanged. In other words, all edges were broken and randomly reconnecting, preserving the degree of every node (Pérez-Bercoff et al. 2011). The distributions of inter-subgenome and intra-subgenome edge counts are shown in Figure 4.

### *Association between recent selective sweeps in* B. rapa *and subgenomes origin*

*B. rapa* genes were divided into those in the regions of selective sweeps detected by SweeD (Pavlidis et al. 2013) in either turnip, toria, Indian sarson, pak choi or Chinese cabbage (vegetable types of *B. rapa*), and those showing no such signatures (Qi et al. 2017, 2021). We tested whether particular subgenomes (posterior probability ≥ 0.95) were unusually likely or unlikely to have experienced a selective sweep using chi-square test. The association plot as shown in Supplemental Fig S7 was visualized using the vcd package version 1.4-4 in R 3.6.0 (Meyer et al. 2006; Zeileis et al. 2007).

## Data Access

The assembled *Crambe hispanica* genome (v1.1) generated in this study has been submitted to the NCBI BioProject database (https://www.ncbi.nlm.nih.gov/bioproject/) under accession number JABFOD000000000. Raw RNA-seq files from *C. hispanica* have been submitted to the NCBI BioProject database under accession number PRJNA475309. The annotation of the *Crambe hispanica* v1.1 genome is available from CoGe (https://genomevolution.org/coge/) under accession id 58014. POInT input files, the inferred ancestral gene orders, POInT models and assumed phylogenetic trees are included in the Supplemental Data and are available from figshare (https://doi.org/10.6084/m9.figshare.12277832) and from the POInT_browse_ portal (http://wgd.statgen.ncsu.edu/).

## Acknowledgements

This project was supported by U.S. National Science Foundation grant NSF-IOS-1339156 (Y.H., M.M., P.P.E., H.A., R.S.A., J.D.W., X.Q., M.B., E.L., J.C.P., and G.C.C.), and by grants NSF-DBI-1743442 and NSF-IOS-2023310 (E.L.). We would like to thank the U.S. Department of Energy Joint Genome Institute and the Brassicaceae Map Alignment Project (BMAP) consortium for allowing us access to the *Sinapis alba* genome. The work conducted by the U.S. Department of Energy Joint Genome Institute, a DOE Office of Science User Facility, is supported by the Office of Science of the U.S. Department of Energy under Contract No. DE-AC02-05CH11231.We would like to thank A. Platts for assistance with our analyses and J. A. Birchler, R. Roberts, J. Thorne and H. Ashrafi for helpful discussions.

## Supplemental Information

**Supplemental Table S1.**
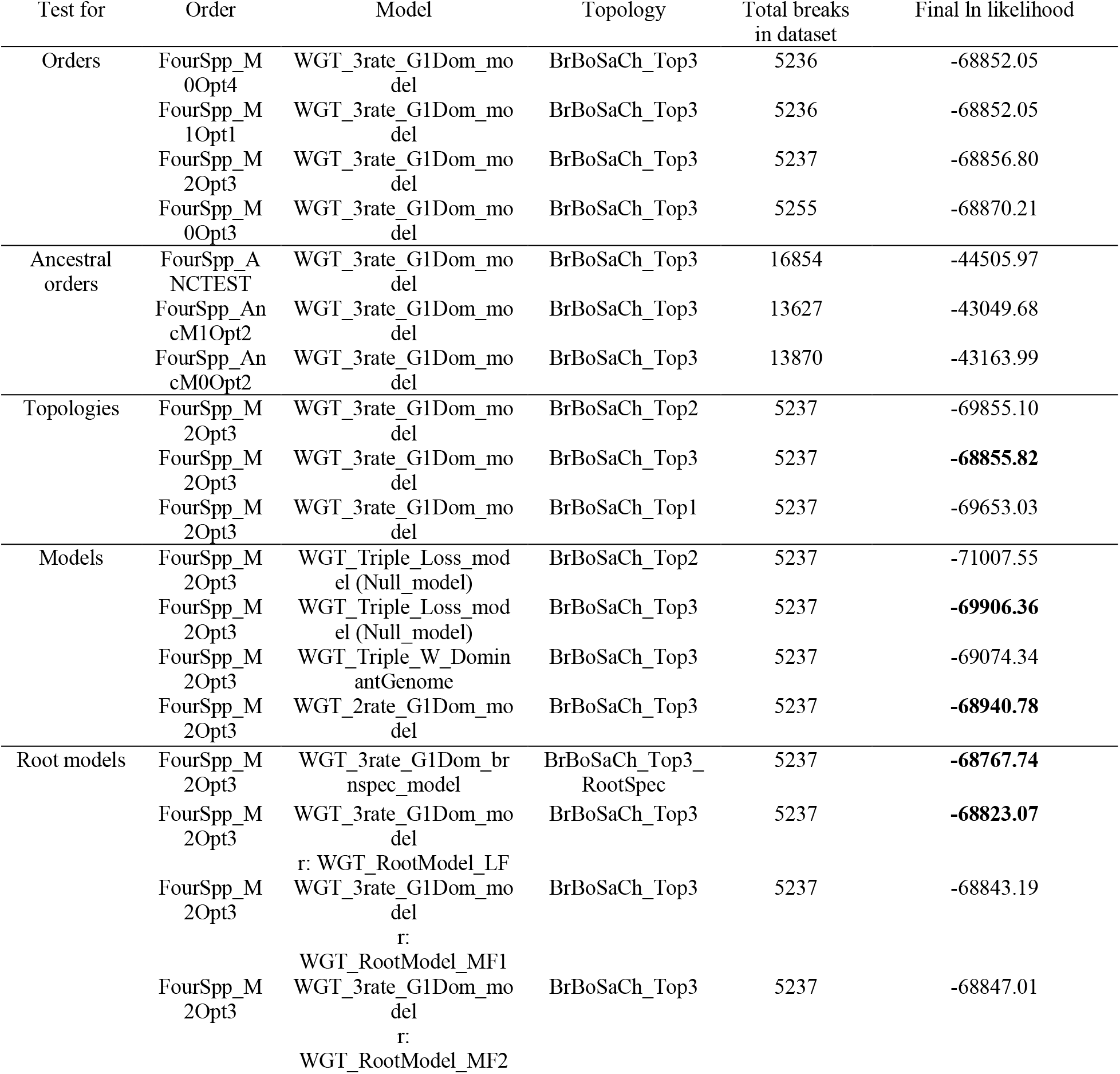
Model optimization and likelihoods.

**Supplemental Fig S1.** Final ln likelihoods of three different topologies of the four species *B. rapa*, *B. oleracea*, *S. alba* and *C. hispanica*.

**Supplemental Fig S2.** Shared synteny blocks across four genomes.

**Supplemental Fig S3.** Species-specific and shared posterior probabilities of all 14,050 loci.

**Supplemental Fig S4.** Selective constraints of retained triplets partitioned into subgenomes.

**Supplemental Fig S5.** PANTHER Biological Processes and Molecular Functions for the Arabidopsis orthologs of genes that returned to single copy at the root branch with FDR ≥ 0.05.

**Supplemental Fig S6.** Number of edges connecting nodes with single copy genes that are from the same subgenome in both protein-protein interaction network and metabolic network.

**Supplemental Fig S7.** *Brassica rapa* subgenome assignment and genes under selective sweep.

## Supplemental Figures

**Supplemental Fig S1.**
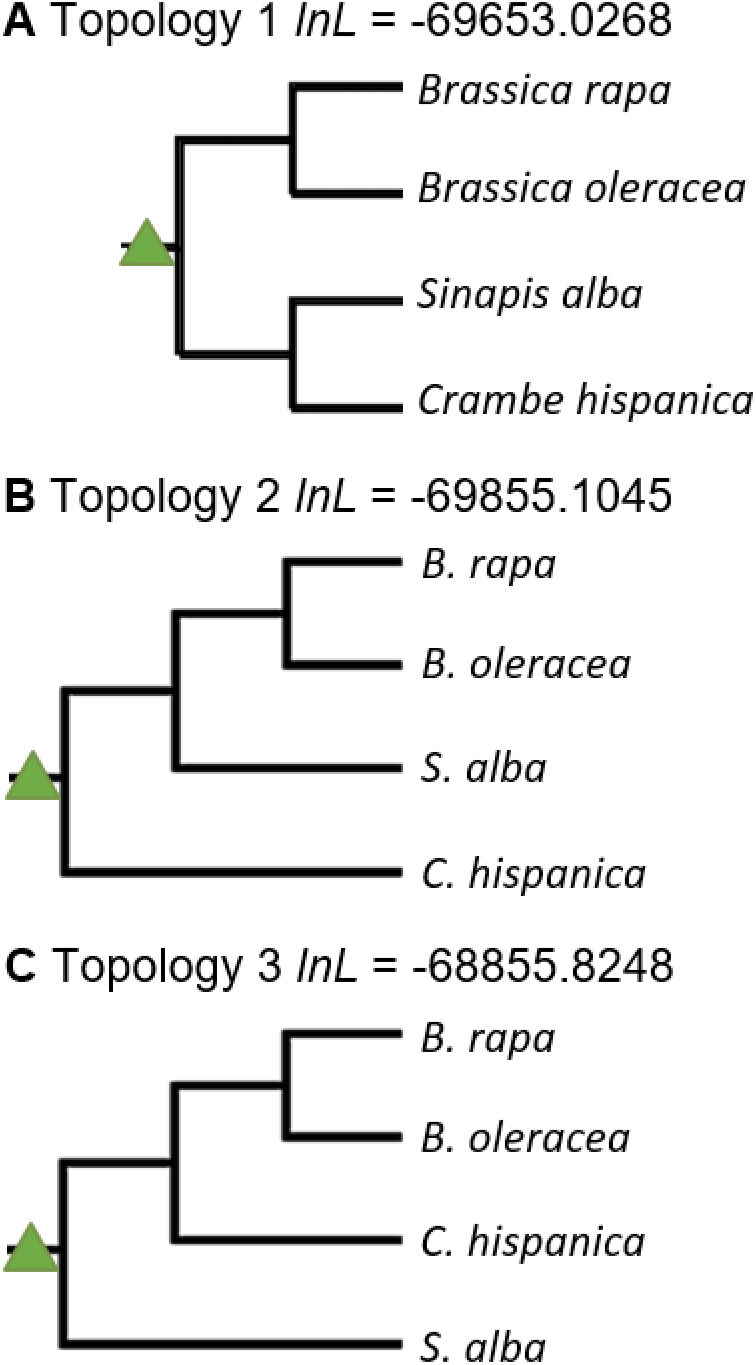
Final ln likelihoods of three different topologies of the four species *B. rapa*, *B. oleracea*, *S. alba* and *C. hispanica*. The triangle indicates shared hexaploidy ancestry.

**Supplemental Fig S2.**
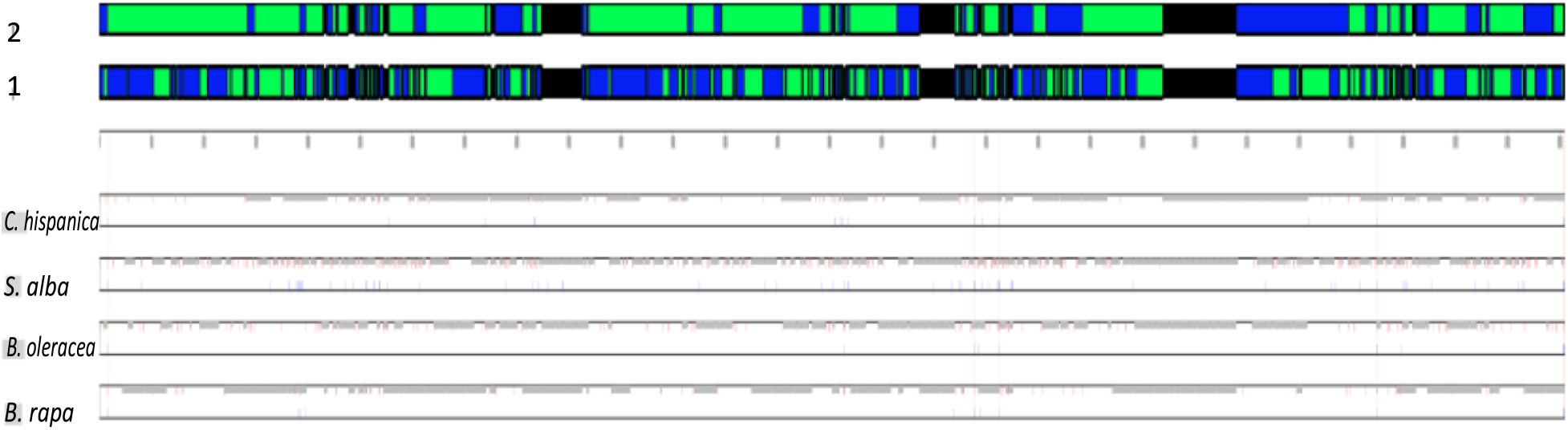
Shared synteny blocks across four genomes. The green and blue blocks indicate shared parental subgenome assignment between at least three (lower blocks) or two (upper blocks) genomes with confidence > 0.85. Each change of color indicates a new block of genes with consistent assignments to the three subgenomes. Black areas indicate a lack of agreement in parental subgenome assignments. The four separate panels below show the POInT subgenome assignment in each species. Red ticks indicate switch in subgenome assignment, grey ticks indicate parental subgenome assignment confidence < 0.85 and blue ticks indicate full synteny breaks in that genome relative to the inferred ancestral order.

**Supplemental Fig S3.**
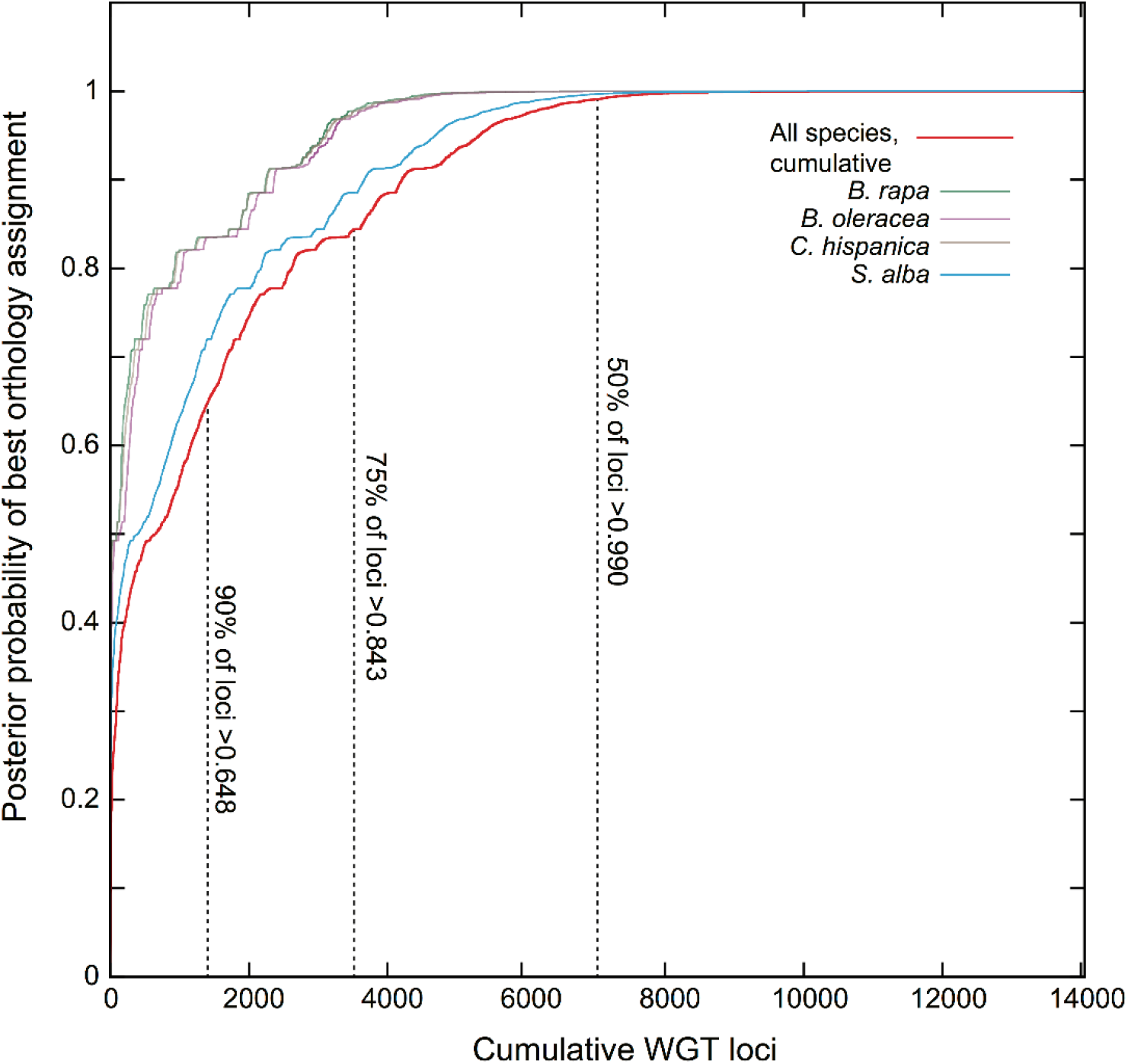
Species-specific and shared posterior probabilities of all 14,050 loci. 50% of the loci have posterior probabilities larger than 0.99, 75% of the loci have posterior probabilities larger than 0.843, 90% of the loci have posterior probabilities larger than 0.648.

**Supplemental Fig S4.**
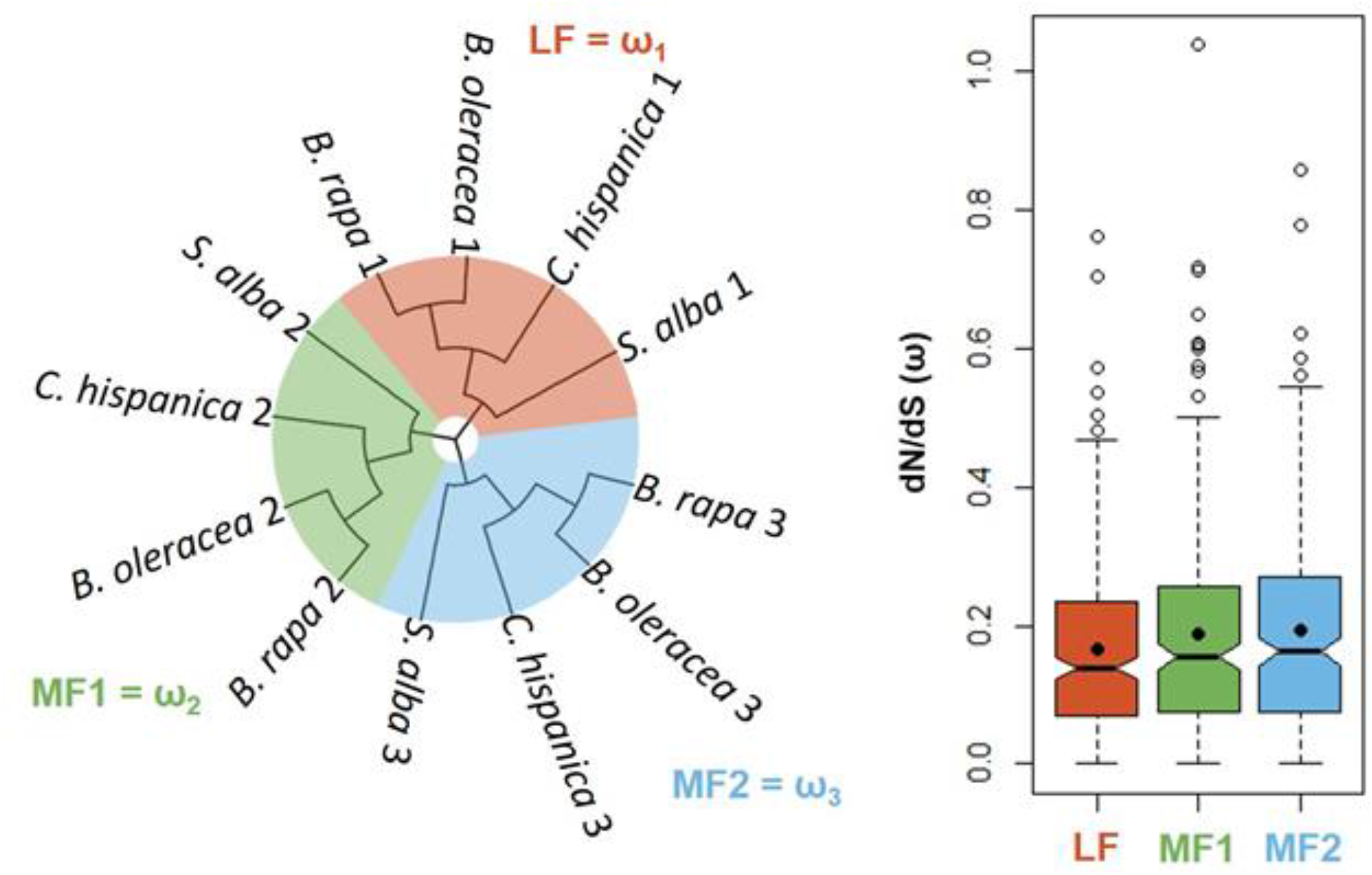
Selective constraints of retained triplets partitioned into subgenomes. As shown in the schematic gene tree, three separate *d_N_/d_S_* values were estimated using codeml for each subtree containing four gene copies that were assigned to the same subgenome in four species. Notched box plots show the distributions of *d_N_/d_S_* for retained copies in each subgenome, LF, MF1 and MF2. The notches show the medians and the 95% confidence intervals. The black dots show the mean values. Pairwise Wilcoxon Rank Sum Tests (Mann and Whitney, 1947) were performed to compare the median selective constraints for retained triplets across subgenomes. LF – MF1: *P* = 0.300 LF – MF2: *P* = 0.079 MF1 – MF2: *P* = 0.516

**Supplemental Fig S5.**
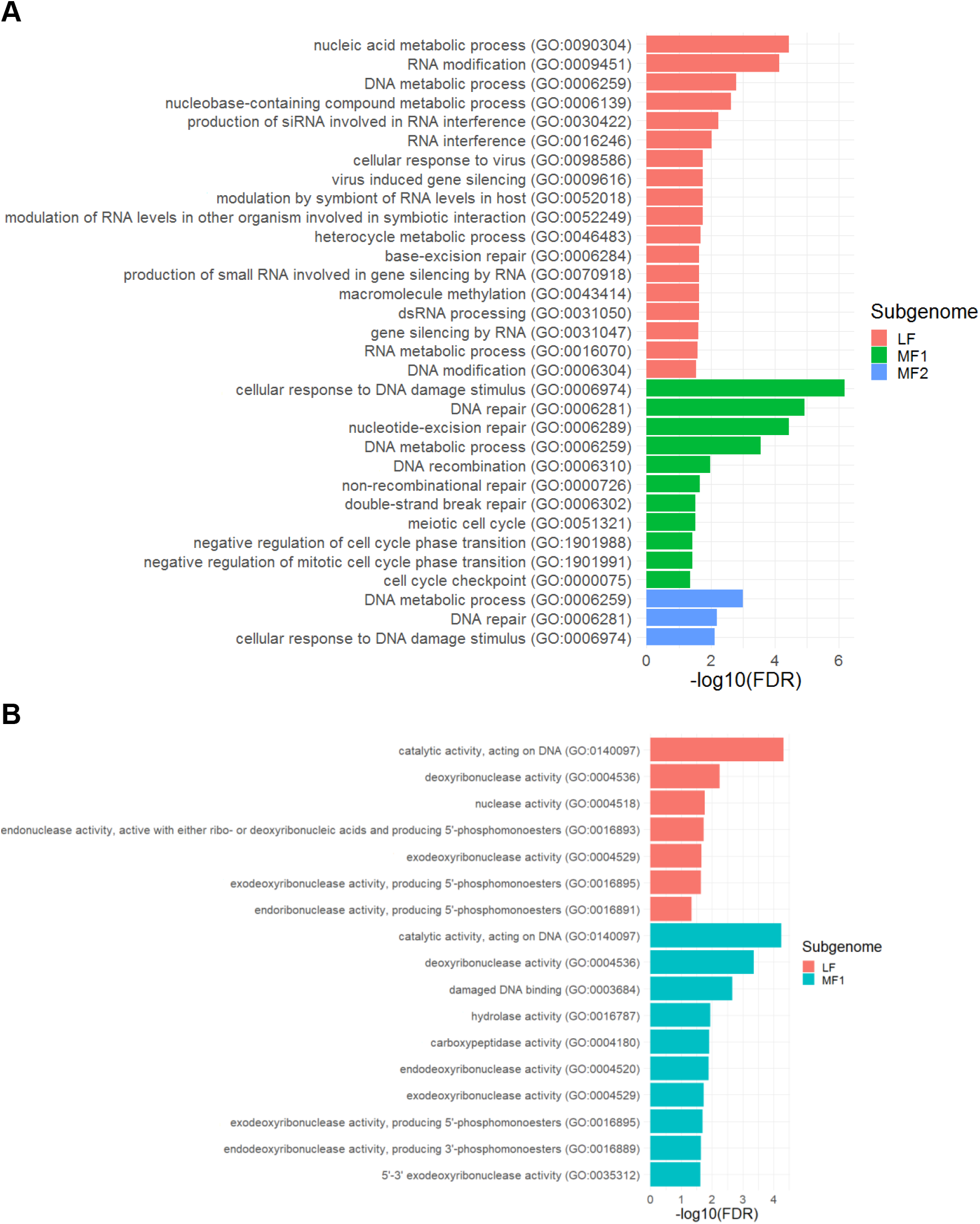
PANTHER Biological Processes (*A*) and Molecular Functions (*B*) for the *Arabidopsis* orthologs of genes that returned to single copy at the root branch with FDR ≥ 0.05. The target lists are single copy genes from three subgenomes LF, MF1 and MF2. The background list was set to be all the retained duplicates and triplets.

**Supplemental Fig S6.**
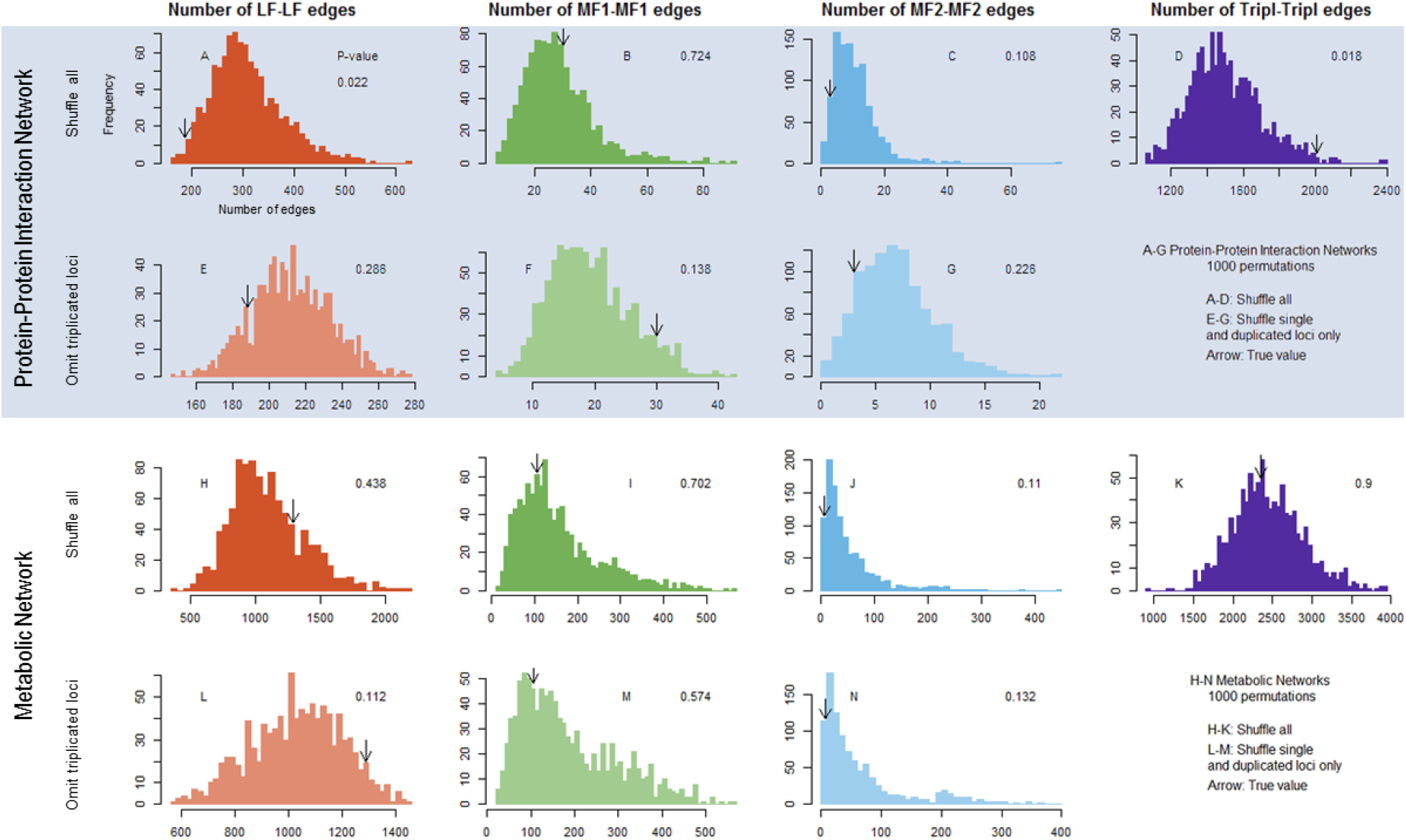
Number of edges connecting nodes with single copy genes that are from the same subgenome in both protein-protein interaction network and metabolic network.

**Supplemental Fig S7.**
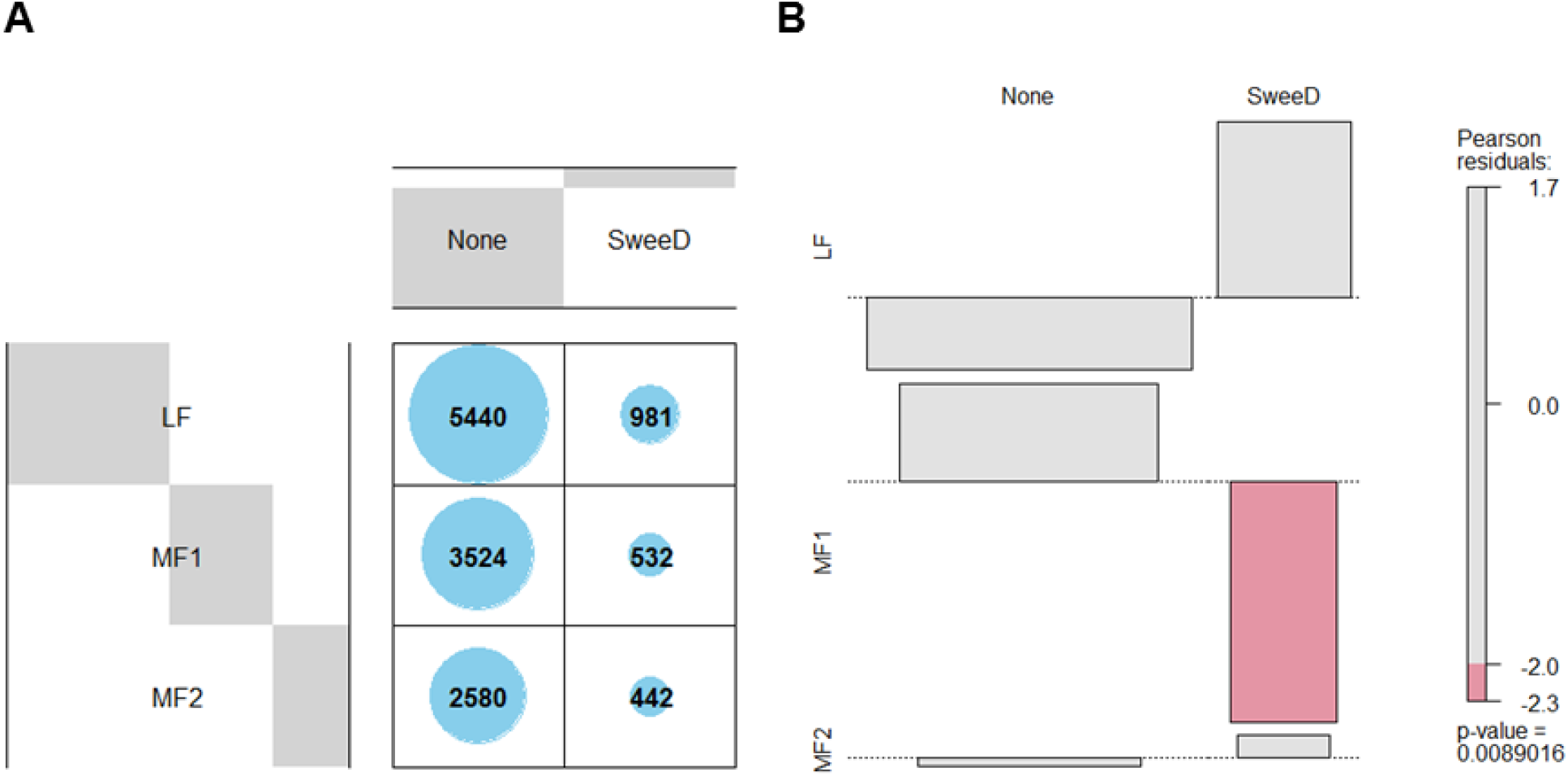
*Brassica rapa* subgenome assignment and genes under selective sweep. (*A*) The number of genes from the three subgenomes (with 0.95 subgenome assignment confidence) versus selective sweeps. (*B*) The association plot based on the contingency table in *A*. The red color in the association plot indicates that the observed value is lower than expected under the random assumption. P-value (0.0089) is from chi-squared test.

